# Derivation of human post-mitotic cardiomyocytes from tetraploid iPSCs

**DOI:** 10.1101/2025.06.23.661005

**Authors:** Ittetsu Nakajima, Mitsuyoshi Shimane, Grace Holmstrom, Yuichiro Miyaoka

## Abstract

Human induced pluripotent stem cell (iPSC)-derived cardiomyocytes (iPS-CMs) have great potential in regenerative medicine. However, iPS-CMs are immature and resemble fetal cardiomyocytes, restricting their application. Although fetal cardiomyocytes in the human heart lose their proliferative potential during maturation and become tetraploid, iPS-CMs cannot replicate this tetraploidization and remain immature. To overcome this problem, we fused diploid iPSCs to establish tetraploid iPSCs and differentiated them into cardiomyocytes (4N-iPS-CMs). We found that 4N-iPS-CMs had more similar gene expression profiles, mitochondrial amounts, contractile impedance, and resistance to a potassium blocker in post-mitotic cardiomyocytes than conventional iPS-CMs. In addition, we successfully generated 4N-iPS-CMs from two individuals to mix two different genetic backgrounds. Thus, we demonstrated a novel strategy for generating human post-mitotic cardiomyocyte-like cells by generating tetraploid iPSCs.

## Introduction

Human induced pluripotent stem cells (iPSCs) have great potential in regenerative medicine^1,2^. However, tissue-specific cells differentiated from iPSCs often exhibit an immature phenotype. For example, most iPSC-derived cardiomyocytes (iPS-CMs) do not become tetraploid^3^, a major hallmark of mature cardiomyocytes^4,5^. Despite tremendous efforts to enhance the maturity of iPS-CMs, no conclusive effective strategies have been developed. Therefore, we shifted our approach to differentiation by first establishing tetraploid iPSC (4N-iPSC) lines and then differentiating them into iPS-CMs to replicate the polyploidy of mature cardiomyocytes in the human heart. We developed a strategy to establish human 4N-iPSCs through cell fusion and found that iPS-CMs derived from 4N-iPSCs (4N-iPS-CMs) were phenotypically similar to post-mitotic cardiomyocytes and were distinct from conventional iPS-CMs. Moreover, we generated 4N-iPSCs derived from two individuals and differentiated them into 4N-iPS-CMs. The resulting 4N-iPSCs and 4N-iPS-CMs will be a powerful platform for life sciences and regenerative medicine.

## Results

### Generation of 4N-iPSCs by fusion of 2N-iPSCs

To generate 4N-iPSCs, we fused diploid iPSCs (2N-iPSCs, WTC11)^6^ with the Hemagluttinating virus of Japan Envelope (HVJ-E) (Figure 1A)^7^. To visualize fused iPSCs, we separately introduced constitutively expressed Venus or mRFP1 into 2N-iPSCs. We mixed and incubated 2×10^5^ of each cell type and HVJ-E for 15 min and plated them. One hour after plating, 20.4% of treated cells were positive for both Venus and mRFP1. After 6 days of culture, 2 colonies positive for Venus and mRFP1 survived (Figure S1A). We selected one of these colonies as our 4N-iPSC cell line (Figure 1B). We first confirmed by flow cytometry that our putative 4N-iPSCs had double the amount of DNA relative to 2N-iPSCs (Figure S1B). DAPI staining of nuclei revealed that the cells had one nucleus rather than multiple nuclei (Figure S1C). We further confirmed, using karyotyping and a CGH analysis, that our fused cells had completely duplicated chromosomes without any microdeletions or insertions (Figures 1C and 1D). Next, we confirmed that these cells expressed the pluripotency markers OCT4, SOX2, and SSEA4 using immunofluorescence staining (Figure S1D). Furthermore, when we transplanted the fused cells into NOD-SCID mice, we observed the formation of teratomas containing all three germ layers (Figure S1E). Based on these results, we conclude that tetraploid iPSCs (4N-iPSCs) were established by cell fusion.

**Figure 1.**
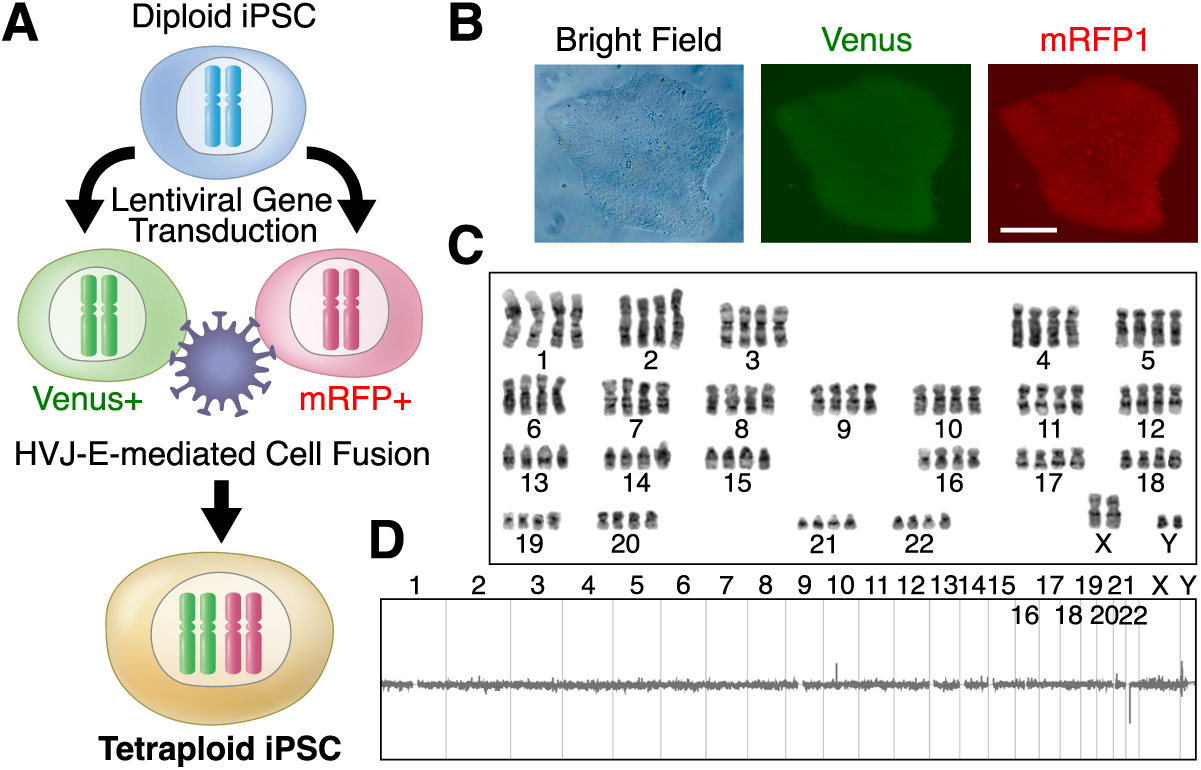
Generation of 4N-iPSCs by fusion of 2N-iPSCs. (A) Venus or mRFP1-positive Japanese male iPSC line, WTC11, was established by lentiviral gene transduction. These cells were fused using Hemagluttinating virus of Japan Envelope (HVJ-E). Fused cells were detected and isolated as Venus and mRFP1 double-positive colonies, followed by single-cell cloning. (B) Fluorescent images of 4N-iPSC colonies. Scale bar, 200 µm. (C) Karyotype of the isolated 4N-iPSCs. Autosomes and sex chromosomes were duplicated without obvious chromosomal abnormality. (D) The comparative genome hybridization (CGH) analysis of the isolated 4N-iPSCs using the original diploid WTC11 as a reference. No chromosomes showed insertions or deletions. See also Figure S1.

### Enhanced sensitivity of 4N-iPSCs to oxidative stress

4N-iPSCs were larger than 2N-iPSCs, as demonstrated by a 1.1-fold increase in diameter (Figure 2A). The doubling time of 4N-iPSCs (22.9 h) was also slower than that of 2N-iPSCs (21.2 h) (Figure 2B). To identify gene expression changes, we performed RNA-seq and found that 618 genes were downregulated, while 159 genes were upregulated in 4N-iPSCs relative to 2N-iPSCs (*P*<0.05, log_2_ |fold change| >1, Figure S2, Table S1). Notably, the expression of metallothionein (MT) family genes was decreased in 4N-iPSCs (Figures 2C and 2D). Since MT proteins bind to divalent metal ions to protect cells from oxidative and metallic stresses, a Gene Ontology enrichment analysis revealed that the top 9 of 10 biological processes predicted to be differentially regulated between 2N and 4N cells were related to the metabolism of metal ions (Figure 2E). Therefore, we treated 2N-and 4N-iPSCs with hydrogen peroxide, copper sulfate, and cadmium chloride to monitor their responses to these cytotoxic reagents and found that 4N-iPSCs were more sensitive to oxidative stress induced by hydrogen peroxide in comparison to 2N-iPSCs (Figure 2F). There was no clear difference in the sensitivity to metallic stress (Figure 2F). Despite these differences, 4N-iPSCs can be regularly cultured and passaged under the same conditions as 2N-iPSCs.

**Figure 2.**
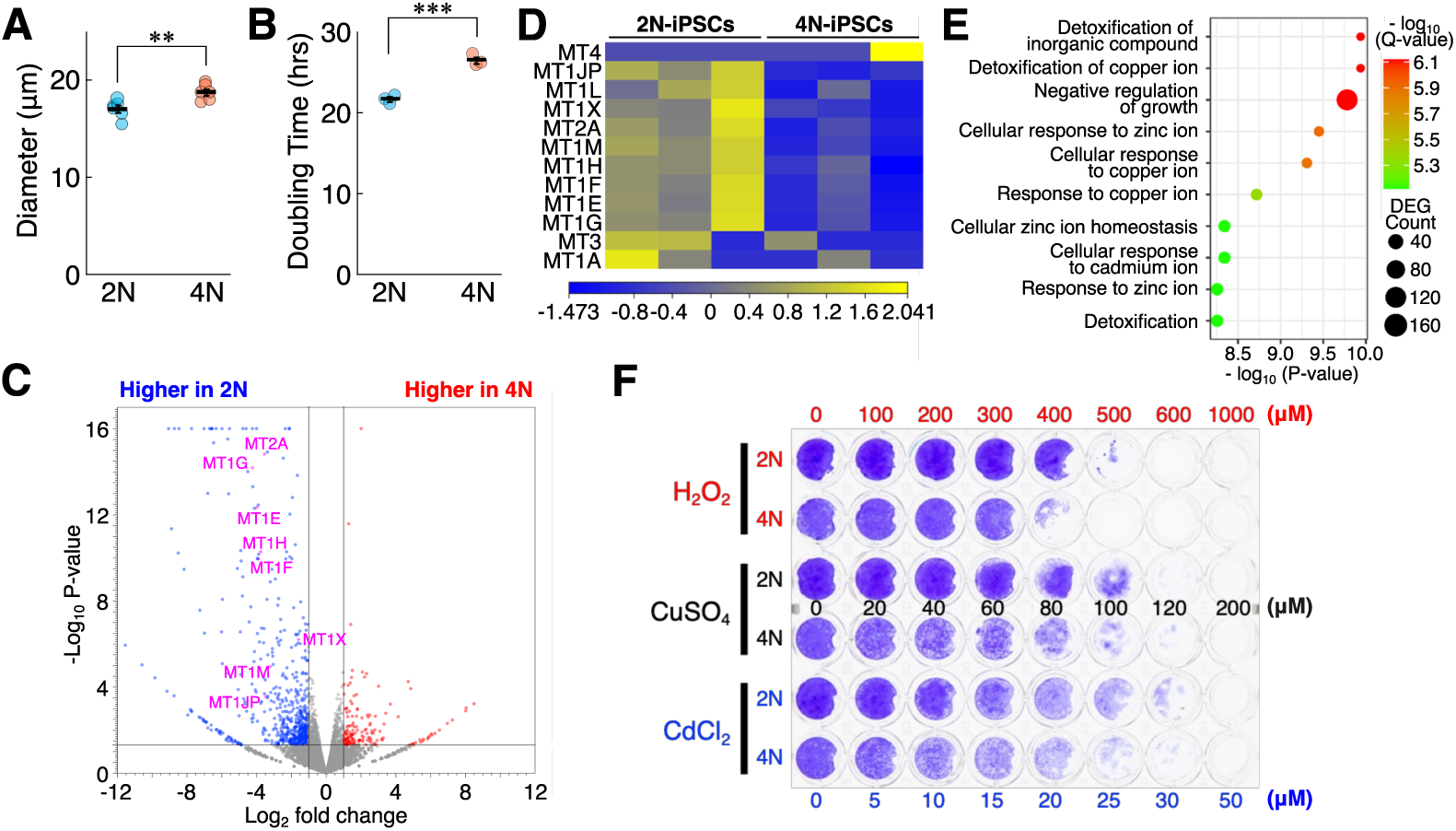
Enhanced sensitivity of 4N-iPSCs to oxidative stress. (A) Diameter of 2N- and 4N-iPSCs (*n*=6). (B) Doubling time of 2N- and 4N-iPSCs (*n*=3). (C) Volcano plot to visualize differentially expressed genes (DEGs) between 2N- and 4N-iPSCs. Red and blue dots represent genes significantly upregulated or downregulated in 4N-iPSCs, respectively (*P*<0.05, |fold-change|>2). Differentially expressed Metallothionein (MT) genes are highlighted in purple. (D) Heatmap representation of the expression of MT genes in 2N- and 4N-iPSCs. (E) The gene ontology (GO) biological process analysis of 2N- and 4N-iPSCs. The top 10 predicted differentially regulated processes between 2N- and 4N-iPSCs are shown. (F) Crystal violet staining showing cell survival of 2N- and 4N-iPSCs in the presence of different concentrations of H2O2, CuSO4, and CdCl2. Error bars, mean ± SEM: \*\**P*<0.01, \*\*\**P*<0.001, two-sided Student’s t test. See also Figure S2.

### Differentiation of 4N-iPSCs into cardiomyocytes

Next, we differentiated 4N-iPSCs into cardiomyocytes using the standard protocol of sequential inhibition of glycogen synthase kinase and Wnt signaling^8^. We found that 4N-iPSCs could be differentiated into beating cardiomyocytes using this protocol, similar to 2N-iPSCs (Videos S1 and S2). The cardiomyocytes were further purified by metabolic selection using lactate^9^. We found that 4N-iPS-CMs were more prone to cell death in lactate, since unlike conventional 2N-iPSC-derived cardiomyocytes (2N-iPS-CMs), they did not survive 6-day selection in lactate. However, 4N-iPS-CMs were successfully purified after 4 days of selection. We next observed organized sarcomeric structures in 4N-iPS-CMs similar to 2N-iPS-CMs by immunofluorescent staining for cardiac troponin T (cTnT) and electron microscopy (Figures 3A and 3B). We further measured the DNA quantity in 4N-iPS-CMs using flow cytometry and confirmed that 4N-iPS-CMs were almost all tetraploid (Figure 3C). Thus, although only approximately 10% of 2N-iPS-CMs became tetraploid (Figure 3C), nearly all 4N-iPS-CMs remained tetraploid after differentiation.

### Decreased expression of mitotic genes in 4N-iPS-CMs

We next characterized 4N-iPS-CMs to determine whether their phenotypes were more similar to mature cardiomyocytes than conventional 2N-iPS-CMs. We first conducted an RNA-seq analysis to examine the global gene expression in 2N- and 4N-iPS-CMs (Table S2). Cell cycle arrest is a major hallmark of mature cardiomyocytes, and we found that genes involved in cell cycle progression, including PCNA, CDC20, PLK1, CCNA2, ANLN, AURKA, AURKB, TOP2A, MKI67, and E2F1, were repressed in 4N-iPS-CMs (Figure 3D). As a result, the Gene Ontology enrichment analysis showed that all the top 10 biological processes predicted to be differentially regulated between 2N- and 4N-iPS-CMs were related to cell cycle regulation (Figure 3E). The expression profiles of genes associated with other cardiomyocyte functions, such as myofibril formation, metabolism, cardiomyocyte subtypes, and electrophysiology, did not show consistent differences between 2N- and 4N-iPS-CMs (Figures S3A-S3D). Thus, 4N-iPS-CMs more closely resemble mature cardiomyocytes in terms of the expression of mitotic genes than 2N-iPS-CMs.

**Figure 3.**
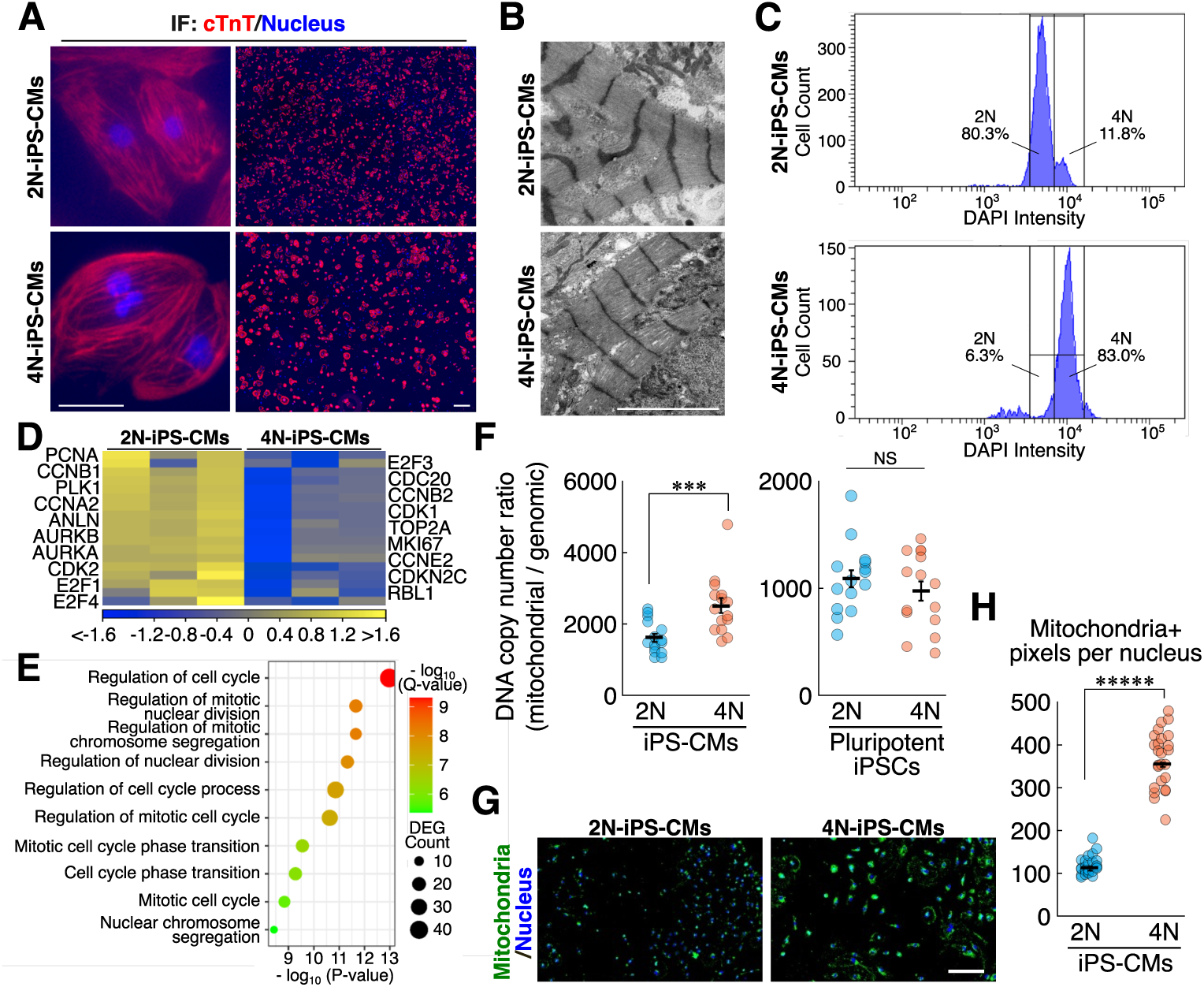
The decreased expression of mitotic genes and increased mitochondria in 4N-iPS-CMs (A) Immunofluorescent staining of cardiac troponin T (cTnT) in 2N- and 4N-iPS-CMs. Scale bars, 50 µm (left) and 200 µm (right). (B) Electron microscopy of the sarcomeric structure. Scale bar, 5 µm. (C) The flow cytometric analysis of the DNA content of 2N- and 4N-iPS-CMs. (D) A heatmap representation of the mitotic gene expression in 2N- and 4N-iPS-CMs. (E) The GO biological process analysis of 2N- and 4N-iPS-CMs. The top 10 predicted differentially regulated processes between 2N- and 4N-iPS-CMs. (F) The copy number ratio of mitochondrial DNA to genomic DNA in 2N- and 4N-iPS-CMs (left) and pluripotent 2N- and 4N-iPSCs (right) quantified by digital PCR (*n*=15). (G) Representative images of mitochondrial staining (green) in 2N- and 4N-iPS-CMs. (H) Quantification of mitochondrial pixel numbers per nucleus in 2N- and 4N-iPS-CMs, as determined by an image analysis (*n*=25). Error bars, mean ± SEM: NS: not significant, *P*>0.05, \*\*\**P*<0.001, \*\*\*\*\**P*<0.00001. See also Figures S3 and S4.

### Increased mitochondria in 4N-iPS-CMs

Another hallmark of mature cardiomyocytes is an increase in mitochondria to meet increased energy requirements^10,11^. To determine whether this increase also occurred in 4N-iPS-CMs, we quantified the copy number ratio of mitochondrial DNA to genomic DNA using digital PCR and found a significantly higher ratio in 4N-iPS-CMs than in 2N-iPS-CMs (Figure 3F). In contrast, the mitochondrial DNA/genomic DNA copy number ratio did not significantly differ between undifferentiated 4N-iPSCs and 2N-iPSCs, indicating that the observed mitochondrial increase was specific to the differentiation process (Figure 3F). We confirmed the increase in mitochondria in 4N-iPS-CMs by visualizing and quantifying them using an image analysis (Figures 3G and 3H). We confirmed an increase in mitochondrial density in 4N-iPS-CMs by electron microscopy (Figure S4). No notable ultrastructural differences were observed between the 2N- and 4N-iPS-CMs (Figures 3B and S4). Therefore, 4N-iPS-CMs more closely resemble mature cardiomyocytes than 2N-iPS-CMs in terms of the increase of mitochondria.

### Increased impedance in 4N-iPS-CMs

Next, to monitor the electrophysiological characteristics of 4N-iPS-CMs, we measured cellular impedance for two weeks after plating cardiomyocytes onto a sensor plate (Figure 4A). 4N-iPS-CMs showed a higher base impedance than 2N-iPS-CMs, suggesting a stronger attachment of the cells to the plate (Figure 4B). There was no significant difference in beat rate between 2N- and 4N-iPS-CMs (Figure S5A), and both cell types exhibited comparable amplitudes for the first six days after plating. However, 4N-iPS-CMs maintained high amplitudes after 6 days, whereas the amplitude in 2N-iPS-CMs peaked at 5 days after plating and then decreased (Figure 4C). This suggests that 4N-iPS-CMs maintain contractile force over time, whereas conventional 2N-iPS-CMs do not. Moreover, while there was no difference in the relaxation velocity (Figure S5B), the upstroke velocity was significantly greater in 4N-iPS-CMs after day 5 (Figure 4D), resulting in different shapes of the impedance peaks between 2N- and 4N-iPS-CMs (Figure 4E). As higher impedance amplitudes and faster upstroke velocities are indicative of mature cardiomyocytes, these results suggest that 4N-iPS-CMs have more mature electrophysiological phenotypes in comparison to 2N-iPS-CMs.

### Increased resistance to a potassium channel blocker in 4N-iPS-CMs

Fetal cardiomyocytes show a higher sensitivity to potassium channel blockers, including terfenadine, and become more resistant to these drugs upon maturation^12^. Thus, we next treated 2N- and 4N-iPS-CMs with different concentrations of terfenadine five days after plating on a sensor plate. 2N-iPS-CMs stopped beating after treatment with 500 and 900 nM terfenadine. In contrast, 4N-iPS-CMs kept beating with 500 and 900 nM terfenadine (Figures 4F, 4G, S5C, and S5D), suggesting that 4N-iPS-CMs were phenotypically more mature than conventional 2N-iPS-CMs.

**Figure 4.**
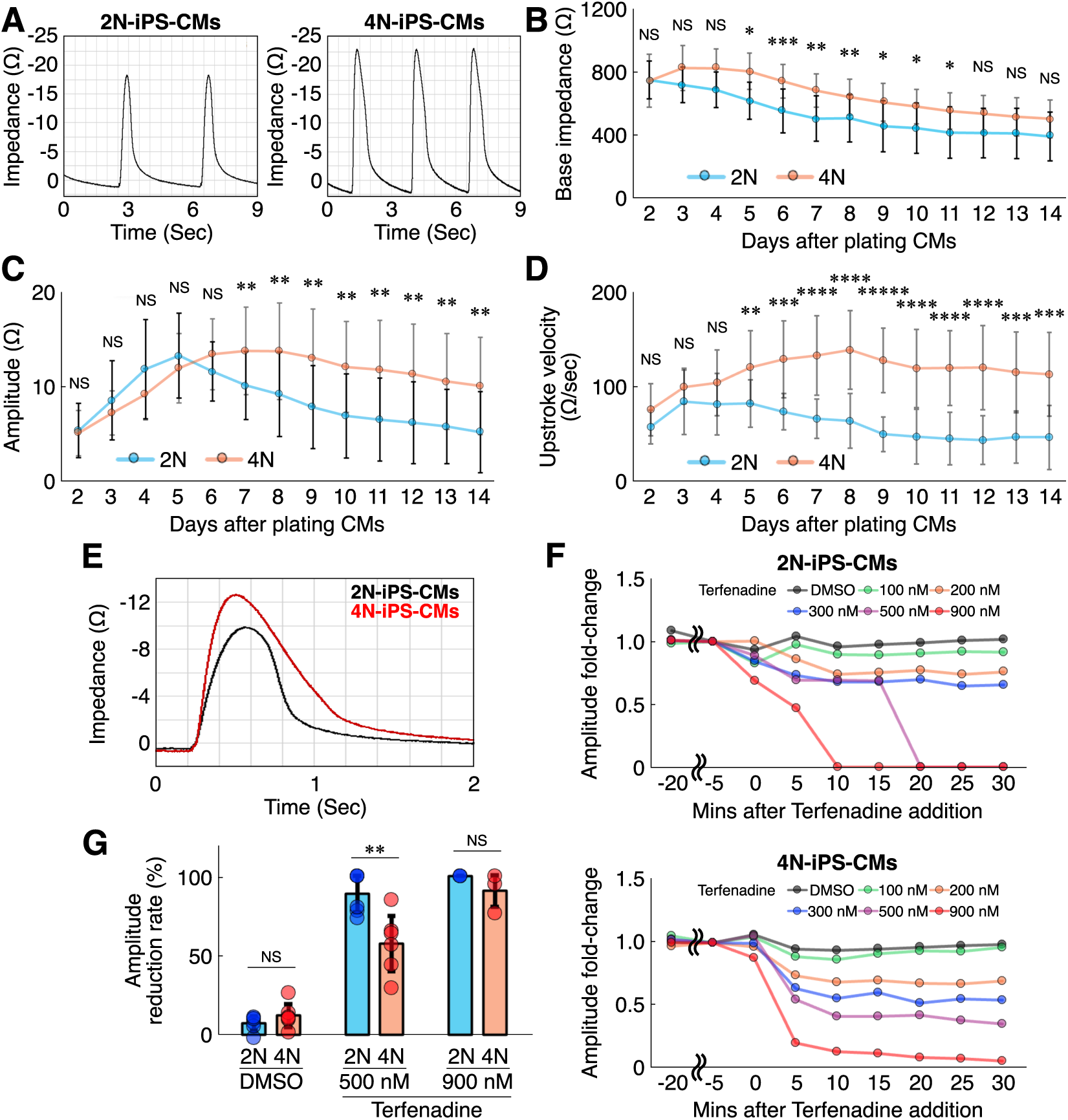
Increased impedance and resistance to a potassium channel blocker in 4N-iPS-CMs. (A) Representative cardiac impedance waveforms of 2N- and 4N-iPS-CMs. (B-D) Base impedance (B), impedance amplitude (C), and upstroke velocity (D) of 2N- and 4N-iPS-CMs over two weeks after plating on a sensor plate (*n*=15 for 2N-iPS-CMs, *n*=11 for 4N-iPS-CMs). (E) Overlayed waveforms of 2N- and 4N-iPS-CMs to highlight the stronger amplitude and faster upstroke velocity of 4N-iPS-CMs. (F) Representative fold-changes of impedance amplitude of 2N- and 4N-iPS-CMs after the addition of different concentrations of Terfenadine. (G) Amplitude reduction rate of 2N- and 4N-iPS-CMs at 30 minutes after the addition of different concentrations of Terfenadine (*n*=6 for DMSO and 500 nM, *n*=3 for 900 nM Terfenadine). Error bars, mean ± SEM: NS: not significant, *P*>0.05, \**P*<0.05, \*\**P*<0.01, \*\*\**P*<0.001, \*\*\*\**P*<0.0001, \*\*\*\*\**P*<0.00001. See also Figure S5.

### Generation of 4N-iPSCs and 4N-iPS-CMs from two different individuals

Thus far, we have generated 4N-iPSCs and 4N-iPS-CMs by fusing 2N-iPSCs from the same individual (WTC11). If we could generate 4N-iPSCs by fusing iPSC lines derived from different individuals and differentiating them into cardiomyocytes, we would be able to analyze the interactions of two different genetic backgrounds in iPSCs and iPS-CMs. To do this, we generated a WTC11 line expressing Venus and the puromycin-resistance gene and an HPS0076 line^13^ expressing mRFP1 and the blasticidin resistance gene. WTC11 was derived from a Japanese male, whereas HPS0076 was derived from a Caucasian female. These cells were fused using HVJ-E and selected for resistance to both puromycin and blasticidin to identify the hybrid 4N-iPSC lines (Figures 5A and S6A). Hybrid 4N-iPSCs contained 4N amounts of DNA (Figure S6B). Karyotyping revealed that 10 of 20 cells maintained double chromosomes, and no marked deletions or insertions were identified by a CGH analysis (Figures 5B and 5C). However, karyotyping of the remaining 10 cells showed chromosomal gain or loss (Figure S6C), suggesting that 4N-iPSCs derived from different individuals may have decreased chromosomal stability in comparison to 4N-iPSCs derived from a single individual. A single nucleotide polymorphism (SNP) analysis by CGH showed that the hybrid 4N-iPSCs had both WTC11 and HPS0076 SNPs on each chromosome (Table S3).

Immunostaining for SOX2, OCT4, and SSEA4 confirmed the expression of the pluripotency genes (Figure S6D). We further conducted RNA-seq of hybrid 4N-iPSCs and HPS0076 and compared them with those of WTC11. A principal component analysis of the RNA-seq data revealed that the gene expression profile of hybrid 4N-iPSCs was located between the profiles of diploid WTC11 and HPS0076 iPSC lines, suggesting that hybrid 4N-iPSCs have mixed transcriptional characteristics, as expected (Figure 5D). Next, we differentiated hybrid 4N-iPSCs into cardiomyocytes and metabolically purified them in the same manner as the WTC11 4N-iPS-CMs. These hybrid 4N-iPS-CMs exhibited organized sarcomere structures and regular beating signatures (Figures 5E and Video S3). We also conducted an RNA-seq analysis of hybrid 4N-iPS-CMs and HPS0076 iPS-CMs and compared them with those of WTC11 2N-iPS-CMs. The gene expression profile of hybrid 4N-iPS-CMs showed features intermediate between those of WTC11 iPS-CMs and HPS0076 iPS-CMs (Figure 5F). These results indicated that 4N-iPSCs and 4N-iPS-CMs can be generated from 2N-iPSCs of different genetic backgrounds.

**Figure 5.**
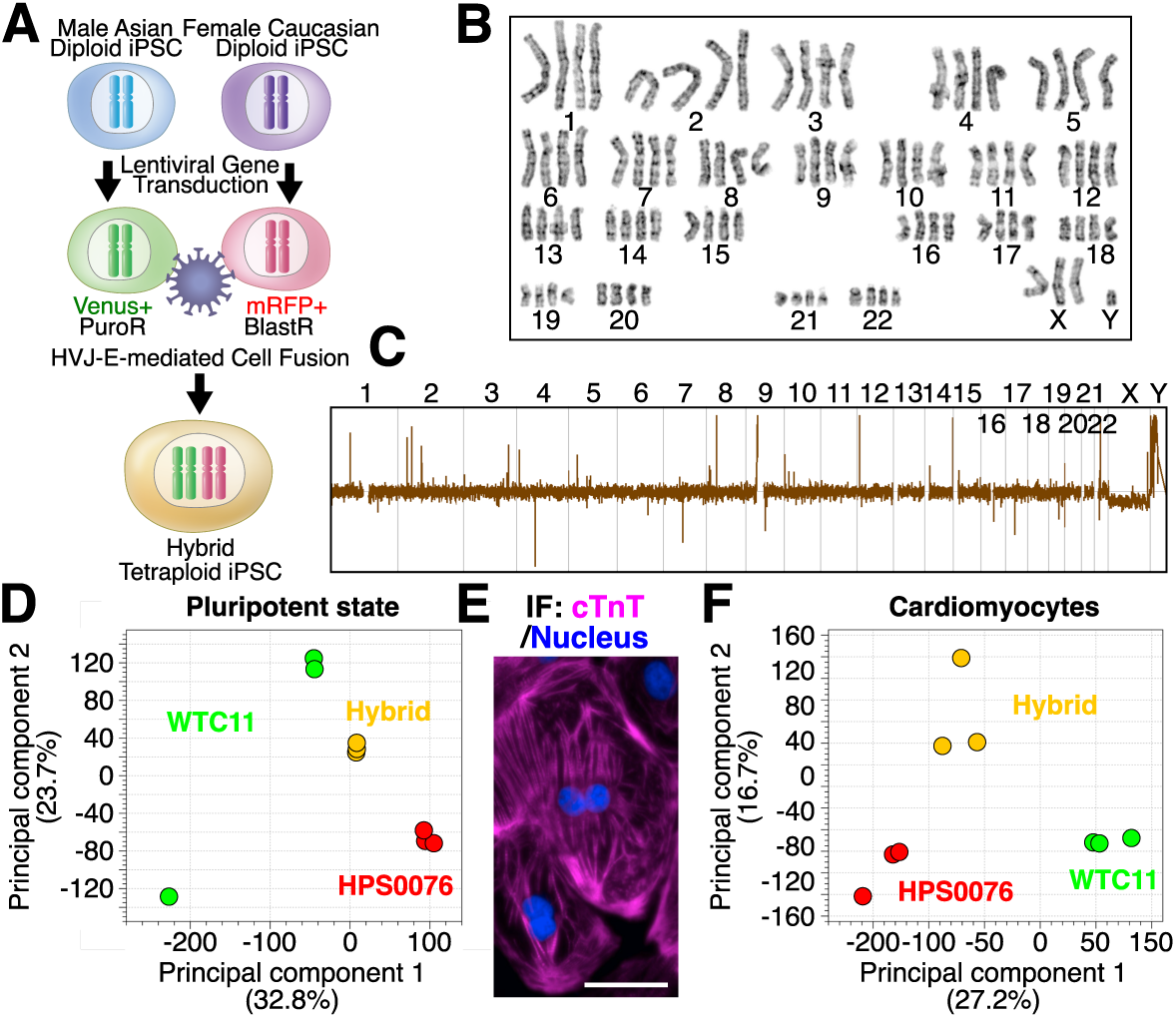
Generation of 4N-iPSCs and 4N-iPS-CMs from two different individuals. (A) The Japanese male iPSC line, WTC11, and the Caucasian female iPSC line, HPS0076, were fused. We established WTC11 and HPS0076 iPSC lines with Venus and puromycin-resistance genes, and mRFP1 and blasticidin-resistance genes, respectively. These cells were fused using HVJ-E. Fused cells were detected and isolated as Venus and mRFP1 double-positive, and puromycin and balsticidin double-resistant colonies, followed by single-cell cloning. (B) The karyotype of the isolated hybrid 4N-iPSCs. Autosomes and sex chromosomes were duplicated without obvious chromosomal abnormalities in this case. (C) The CGH analysis of the isolated hybrid 4N-iPSCs using a standard male diploid iPSCs as a reference. No chromosomes showed insertions or deletions. (D) The principal component analysis (PCA) of WTC11 2N-iPSCs, HPS0076 2N-iPSCs, and hybrid 4N-iPSCs. Hybrid 4N-iPSCs were positioned between the two 2N-iPSCs. (E) Immunostaining of cardiac troponin T (cTnT, magenta) to confirm successful cardiomyocyte differentiation of hybrid 4N-iPSCs. (F) PCA of WTC11 2N-iPS-CMs, HPS0076 2N-iPS-CMs and hybrid 4N-iPSCs. Hybrid 4N-iPS-CMs were positioned between the two diploid cardiomyocytes. See also Figure S6.

## Discussion

Tremendous efforts have been made to improve the maturity of iPSC-derived cardiomyocytes, including optimization of the culture medium, addition of small molecules or cytokines, genome editing, electrical stimulation, 3D-culture, and transplantation into the mouse heart^14^. However, none of these approaches have successfully induced polyploidization in cardiomyocytes, a major hallmark of maturation. We overcame this limitation by first generating tetraploid iPS cells (4N-iPSCs) by cell fusion (Figure 1). While the fusion of pluripotent stem cells with other cell types has been demonstrated previously, the main focus of these studies was on reprogramming differentiated cells into a pluripotent state, and the biological and therapeutic potential of tetraploid pluripotent cells has not been thoroughly investigated^15,16^. Generation of triploid embryonic stem (ES) cells by the fusion of haploid and diploid ES cells has also been reported^17^, but very few triploid cells exist in the body under physiological conditions. In this study, we demonstrated that the fusion of 2N-iPSCs can be used to generate 4N-iPSCs that can be differentiated to produce 4N-iPS-CMs that are phenotypically distinct from conventional 2N-iPS-CMs and more closely resemble mature cardiomyocytes (Figures 3 and 4).

Over 80% of cardiomyocytes in humans become tetraploid during maturation, but the biological significance of this polyploidy remains unknown^4^. One possibility is that diploid cardiomyocytes may serve as heart stem cells and proliferate, whereas tetraploid cardiomyocytes become post-mitotic^18^. Our observation that tetraploidy is associated with the repression of cell cycle genes in iPS-CMs is consistent with this speculation (Figures 3D and 3E). Moreover, 4N-iPS-CMs exhibited increases in mitochondria and more mature electrophysiological phenotypes than 2N-iPS-CMs (Figures 3F-3H). Despite these distinct phenotypes, we did not observe significant differences in the expression of genes associated with sarcomeres, metabolism, or ion channels between the 2N- and 4N-iPS-CMs (Figure S3), suggesting that tetraploidy itself might enhance the maturation of cardiomyocytes. Furthermore, we observed elevated resistance to terfenadine in 4N-iPS-CMs compared to 2N-iPS-CMs (Figures 4F and 4G), possibly due to the greater maturity of 4N-iPS-CMs. The immaturity of iPS-CMs has become an issue of increasing concern in drug screening and cell therapy. Cells expressing the human Ether-a-go-go-Related-Gene (hERG) are currently used to assess the cardiotoxicity of drug candidates^19^, iPS-CMs are expected to replace this artificial system. Our study indicates that 4N-iPS-CMs may be more useful for cardiotoxic assessment than hERG-expressing cells and conventional 2N-iPS-CMs.

In this study, we focused on cardiomyocytes. However, there are various other polyploid cell types in the body, including hepatocytes, skeletal muscle cells, and osteoclasts. It is tempting to speculate that the maturation of these cell types might also be enhanced if they are differentiated from 4N-iPSCs. Moreover, comparisons of cell types differentiated from 2N- and 4N-iPSCs may help us understand the biological significance of polyploidy. Indeed, we were able to identify that the MT family genes were downregulated in undifferentiated 4N-iPSCs relative to 2N-iPSCs (Figures 2C and 2D). This led us to conclude that 4N-iPSCs are more prone to oxidative stress (Figure 2F). Interestingly, a method known as tetraploid complementation has been used to examine the pluripotency of cells in mice. In tetraploid complementation, diploid cells of interest are injected into a tetraploid blastocyst generated by electrofusion of two blastomeres. The tetraploid cells in these chimeric blastocysts were eventually eliminated from the embryo and contributed only to extraembryonic tissues. If the injected diploid cells are truly pluripotent, mouse embryos derived solely from the injected diploid cells develop, demonstrating the pluripotency of the injected cells^20,21^. While the mechanism for the elimination of tetraploid cells is unknown, our finding that tetraploid pluripotent stem cells have increased sensitivity to oxidative stress may be key to elucidating this mechanism. Thus, comparison of various cell types differentiated from 2N- and 4N-iPSCs will be a powerful tool for studying the significance of ploidy differences.

Finally, we demonstrated that 4N-iPSCs can be generated by fusing 2N-iPSCs derived from different individuals, although the genomic stability may be reduced in this case (Figures 5 and S6). This method of introducing two different genetic backgrounds in the same cells may be useful, particularly for physiologically polyploid cells, such as cardiomyocytes. For example, it may be possible to at least partially model maternal and paternal genetic interactions in cardiomyocytes and hepatocytes of progeny in vitro by the fusion of maternal and paternal iPSCs. We propose that 4N-iPSCs provide a powerful platform for studying the significance of ploidy in various cell types and 4N-iPS-CMs, in particular, as a promising tool for drug development and regenerative medicine.

## Resource availability

### Lead contact

Further information and requests for resources and reagents should be directed to and will be fulfilled by the lead contact, Yuichiro Miyaoka.

### Materials availability

- Plasmids generated in this study have been deposited to [Addgene, name and catalog number or unique identifier].
- All iPSC lines generated in this study will be made available upon request, but we may require a payment and/or a completed material transfer agreement.
- All reagents and materials used in this manuscript are available upon request or prepared for availability from commercial sources

### Data and code availability

- RNA-seq data have been deposited at the DDBJ Sequence Read Archive: accession number and are publicly available as of the date of publication.
- The original codes for quantification of mitochondria amount and counting nuclei by ImageJ software are available in this paper’s supplemental information.
- Any additional information required to reanalyze the data reported in this paper is available from the lead contact upon request.

## Acknowledgments

We thank Y. Nishito, Y. Iguchi, K. Endo, K. Mikami, E. Kawakami, J. Horiuchi, and T. Yagi (Tokyo Metropolitan Institute of Medical Science) for their technical help with the CGH analysis, tissue sectioning, electron microscopy analysis, and critical reading of the manuscript. We thank all lab members for their helpful discussions. Funding: This work was supported by the following grants: The Japan Society for the Promotion of Science (JSPS) Grant-in-Aid for Challenging Research (Pioneering) (24K21954) to Y.M., JSPS Grant-in-Aid for Scientific Research (B) (24K02028) to Y.M., Japan Cardiovascular Research Foundation, The Bayer Scholarship for Cardiovascular Research to Y.M., SENSHIN Medical Research Foundation Grant to Y.M., TERUMO Life Science Foundation Research Grant to Y.M., Japan Science and Technology Agency (JST) SPRING (JPMJSP2120) to I.N., and JSPS Grant-in-Aid for JSPS Fellows (24KJ0995) to I.N.

## Author contributions

Conceptualization, I.N. and Y.M.; methodology, I.N., M.S., G.H., and Y.M.; Investigation, I.N., M.S.., G.H., and Y.M.; writing—original draft, I.N. and Y.M.; writing—review & editing, I.N., M.S., G.H., and Y.M.; funding acquisition, I.N. and Y.M.; resources, I.N., M.S., G.H., and Y.M.; supervision, Y.M.

## STAR Methods

### Cell lines

HEK293T (RRID: CVCL_0063) cells were maintained in Dulbecco’s modified Eagle medium (DMEM) with high glucose, sodium pyruvate, and L-glutamine (Thermo Fisher Scientific, 11965118) supplemented with 10% FBS and 1% Penicillin-Streptomycin (P/S, Sigma-Aldrich, P4333). For passaging, cells were detached using 0.25% trypsin-EDTA (Gibco, 2520056).

WTC11 (RRID: CVCL_Y803) and HPS0076 (RRID: CVCL_K092) iPSCs were maintained on Matrigel Matrix (Corning, 356231) in mTeSR Plus medium (STEMCELL Technologies, ST-100-0276) supplemented with 1% P/S, and the medium was changed every two days. When cell density was low, 10 μM Y-27632, a Rho-associated kinase (ROCK) inhibitor (Ri, FUJIFILM Wako Pure Chemical, 034-24024), was added to enhance cell survival. When the cells reached 80-90% confluency, they were detached using Accutase (Nacalai Tesque, 12679-54) and passaged.

4N-iPSCs were maintained under the same culture condition to 2N-iPSCs.

### Production and infection of lentivirus particles

The lentiviral plasmids CSII-EF1α-VENUS-IRES2-PuroR, CSII-EF1α-mRFP1-IRES2-BsdR, CSII-EF1α-H2BC11-VENUS-IRES2-PuroR, and CSII-EF1α-H2BC11-mRFP1-IRES2-BsdR were constructed by replacing the Green Fluorescence Protein (GFP) and/or puromycin resistance (PuroR) gene cDNAs of CSII-EF-GFP-IRES-PuroR (RDB12868, RIKEN BRC) with VENUS, mRFP1, and blasticidin resistance (BsdR) gene cDNAs or by inserting histone H2BC11 cDNA. Twenty-four hours before plasmid transfection, HEK293T cells were plated on two 10-cm dishes each containing 6 × 10^6^ cells in DMEM supplemented with 10% FBS and 1% P/S. After 18 h, the culture medium was replaced with a fresh medium. Six hours later, 8.1 μg of lentiviral plasmid, 5.4 μg of pMDLg/pRRE (Addgene #12251) and 4.5 μg of pCMV-VSV-G-RSV-Rev (RDB04393, RIKEN BRC) were mixed in 1.8 ml of Opti-MEM (Gibco) with 54 μl of 1 mg/ml PEI MAX (Polysciences, Inc.) per dish and incubated for 25 min at room temperature. The mixture was dripped onto a 10-cm dish for transfection. Sixteen hours after transfection, the culture medium was removed and 12 ml of DMEM supplemented with 10% FBS and 1% P/S containing 10 mM HEPES buffer (Gibco) was added. Fifty-five hours later, the culture medium containing lentivirus particles was collected from two 10-cm dishes into a 50-ml conical tube. The dishes were rinsed with 2 ml of phosphate-buffered saline (PBS), and the PBS was collected into the same 50-ml conical tube. The collected medium was centrifuged at 1200 × *g* for 20 min at 4°C. The supernatant was transferred into a new 50-ml conical tube by passing through a 0.45 μm filter (Sartorius) to remove cellular debris. Nine milliliters of Lenti-X™ concentrator (Takara Bio) was added to the filtrate and mixed by conversion. The mixture was incubated at 4°C for 30 min and centrifuged at 1500 × *g* at 4°C for 45 min. The supernatant was removed and the pellet was dissolved in 150 μl of mTeSR Plus with P/S. The viral suspension was transferred into a cryotube (Greiner) and stored at -80°C. To infect lentivirus into iPSCs, 1.25 × 10^5^ iPSCs in mTeSR Plus with Ri and P/S were plated into a well of a 12-well plate. The next day, the culture medium was changed to 1 ml of mTeSR Plus with P/S without Ri, to dissolve the lentivirus particles. The medium was then changed with 1 ml of mTeSR Plus with P/S without lentivirus particles every two days. When the cells reached 80–90% confluency, they were washed three times with 1 ml of PBS to remove the lentivirus particles. The washed cells were detached using Accutase and plated into a new well with 1 ml of mTeSR Plus with Ri, P/S and 0.5 μg/ml puromycin or 10 μg/ml blasticidin to select infected cells. To isolate iPSC clones, antibiotic-selected cells were plated on 96-well plates at 3 or 5 cells per well. When the wells contained only one colony, these colonies expanded as iPSC clones.

### Fusion of iPSCs

Hemagglutinating Virus of Japan Envelope-mediated cell fusion was conducted using the GenomOne-CF kit (Ishihara Sangyo). iPSCs were cultured in a 6-well plate until they reached 70-80% confluency and detached using Accutase. We collected 2 × 10^5^ iPSC lines into a 1.5 ml microtube and centrifuged at room temperature at 200 × *g* for 3 min. The supernatant was removed and 50 μl of cell fusion buffer (Ishihara Sangyo) was added to the pellet and mixed gently. The cell suspension was transferred to a cold 1.5 ml microtube with 1 μl of HVJ-E (Ishihara Sangyo) and mixed gently. The mixture was incubated for 5 min on ice and centrifuged at 200 × *g* at 4°C for 5 min. The cells were incubated for 15 min at 37°C without removing the supernatant. After incubation, the pellet was tapped and dissolved in 10 ml of mTeSR Plus containing Ri and P/S. One hundred microliters of cell suspension per well was dispensed into a 96-well plate. The medium was replaced with fresh mTeSR Plus with Ri and P/S every two days. Cells positive for both Venus and mRFP1 fluorescence were passaged into a 12-well plate with 1 ml of mTeSR Plus containing Ri, P/S, 0.5 μg/ml puromycin, and 10 μg/ml blasticidin to select the fused cells. The ratio of Venus and/or mRFP1 single- or double-positive cells was calculated using the ImageJ software program.

### Cell cycle assay

Cells were detached from the wells using Accutase and resuspended in PBS. We collected 5 × 10^5^ cells in a 1.5 ml-conical tube and centrifuged at 300 × *g* at room temperature for 3 min. After removing the supernatant, the cells were washed with 1 ml of PBS and centrifuged again. After removing the supernatant, 1 ml of 4% paraformaldehyde (PFA) was added to the cell pellet and incubated at room temperature for 20 min. The cell suspension was centrifuged at 300 × *g* at room temperature for 3 min and the supernatant was removed. The cells were washed with 1 ml of PBS and centrifuged again. The cell pellet was resuspended in 500 μl of PBS. Five microliters of cell cycle assay solution blue (Dojindo) was added to the cell suspension and incubated for 15 min at 37°C. The cell suspension was passed through a 35-μm filter and stored on ice. Fluorescence at 450/50 nm was measured using a BD LSRFortessa™ X-20 flow cytometer (Becton Dickinson).

### Observation of nuclei of iPSCs

iPSCs were plated in mTeSR Plus with Ri and P/S in a 24-well plate at 2 × 10^4^ cells and 1 × 10^4^ cells per well for 2N- and 4N-iPSCs, respectively. The next day, the culture medium was removed and the wells were washed with PBS three times. The cells were then stained with 10 μg/ml Hoechst 33342 for 10 min and washed with PBS three times. Images were obtained using a fluorescence microscope BZ-X810 (Keyence) and analyzed using the ImageJ software program.

### Karyotyping

A G-band karyotyping analysis of the cell lines was performed by Nihon Gene Research Laboratories.

### CGH analysis

Genomic DNA was extracted from the cells using the DNeasy Blood & Tissue Kit (Qiagen), and a CGH analysis was performed using the SurePrint G3 Human CGH 4x180K Microarray or SurePrint G3 CGH & CGH+SNP Microarray.

### Fluorescent immunostaining

Wells were washed three times with PBS. Then, 4% paraformaldehyde (PFA) was added to the wells and incubated for 20 min at room temperature for fixation. After washing the wells three times with PBS, the cells were incubated and permeabilized with PBS containing 3% normal goat serum (Jackson ImmunoResearch LABORATORIES Inc.) and 0.2% Triton X-100 (Wako) for 30 min at room temperature. Primary antibodies against SOX2 (Abcam, ab97959, 1:500), OCT4 (Abcam, ab97959, 1:500), SSEA4 (Abcam, ab16287, 1:500), and Cardiac Troponin T (Invitrogen, MA5-12960, 1:200) diluted with PBS containing 3% normal goat serum and 0.2% Triton X-100 were added to the cells and incubated at 4°C overnight. After washing the wells with PBS three times, secondary antibodies (Invitrogen, Alexa Fluor 647, A21235, A21244, Alexa Fluor 568, A11004, 1:1000) diluted in PBS containing 3% normal goat serum and 0.2% Triton X-100 were added to the cells and incubated at room temperature for 1 h. After washing the wells three times with PBS, 1 μg/ml Hoechst 33342 diluted with PBS was added to the wells and incubated at room temperature for 10 min to stain the nuclei. After washing three times with PBS, images were obtained using a fluorescence microscope BZ-X810 (Keyence).

### Teratoma formation and confirmation of three germ layers

iPSCs were cultured in a well of a 6-well plate and detached using 500 μl of 0.5 mM EDTA (Nacalai Tesque). The cells were resuspended in 1.5 ml of mTeSR Plus with 10 μM Y-27632. A cell lifter (Greiner) was used to scrape the cells while keeping the cell clumps. One million cells were collected in a 1.5 ml microtube and centrifuged at 200 × *g* at room temperature for 3 min. The supernatant was removed, and a mixture of 50 μl of GFR Matrigel Phenol Red Free and 50 μl of mTeSR Plus with 10 μM Y-27632 was added to gently mix the cells. The cell suspension was aspirated using a 1 ml syringe (TERUMO) with an 18 G needle (TERUMO) and stored on ice until transplantation. NOD.Cg-Prkdcscid/J (RRID: IMSR_JAX:001303, Jackson Laboratory) mice were anesthetized with 2% isoflurane (VIATRIS). A 1-cm incision was made in the abdominal region, and the epididymal fat pad was carefully pulled out along with the testis. The cell suspension was injected using a syringe and maintained for 10 s to avoid backflow. The testis was pushed back to its original location and the incision was sutured. After 8 weeks, teratomas were excised from the mice, washed with PBS, and fixed with 4% PFA at 4°C overnight. The fixed tissue was washed with water for 10 min and incubated overnight in 20% sucrose in PBS at 4°C. The tissue was incubated overnight in 30% sucrose in PBS at 4°C. The tissue was frozen and sectioned at a thickness of 6 μm. The sections were stained with hematoxylin and eosin to analyze tissue structures.

### Measurement of diameter of iPSCs

The diameter of the iPSCs on a Countess Cell Counting Chamber Slide (Thermo Fisher Scientific) was automatically calculated using Countess II (Thermo Fisher Scientific). Each well of the chamber slide contained >50 cells.

### Calculation of doubling time of iPSCs

Diploid iPSCs (1 × 10^5^ cells) or tetraploid iPSCs (5 × 10^4^ cells) were plated into a well of a 6-well plate (day 0) and cultured for 5 days with changing medium on days 2 and 4. Cells were detached using Accutase, and the number of cells was counted on day 5. These steps were repeated six times (for 30 days) and the doubling time was calculated by dividing the total cell count by the number of cultures.

### RNA-seq

Total RNA was extracted, and DNA was digested using the RNeasy Mini Kit (Qiagen, 74104). The RNA samples were sent to Azenta LIFE SCIENCES. Libraries were prepared using the MGIEasy RNA Directional Library Prep Set V2.0. Sequencing was performed using a DNBSEQ (MGI Tech). For a gene expression analysis, reads were mapped to the GRch38/hg38 reference genome, and the following analysis was conducted using the CLC Genomics Workbench (Qiagen).

### Survival assay of iPSCs to oxidative and metallic stress

We cultured 2N- and 4N-iPSCs in mTeSR Plus in a 96-well plate until they reached confluence. When the cells reached confluence, the medium was changed to Essential 8 (Thermo Fisher Scientific) with different concentrations of H2O2, CuSO4, and CdCl_2_. The next day, the medium was removed, and 30 μl/well of ice-cold 100% methanol was added to fix the cells for 10 min. Methanol was removed, and 30 μl/well of 0.5% crystal violet staining solution (500 mg crystal violet in 25 ml of 100% methanol and 75 ml of water) was added to stain the cells for 10 min. The staining solution was removed, and the plate was washed with running tap water for 10 min. The plates were dried for observation.

### Cardiomyocyte differentiation of human iPSCs

We followed a previously reported protocol for cardiomyocyte differentiation (*8*, *22*^22^. Briefly, 2 × 10^5^ diploid or 1.5 × 10^5^ tetraploid iPSCs in mTeSR Plus with Ri and P/S were plated into a 12-well plate. The next day, the medium was replaced with mTeSR Plus without Ri and P/S. The next day, the medium was changed to RPMI1640 with B-27 minus insulin (Thermo Fisher) and 6 μM CHIR99021 (ChemScene) (day 0). On day 2, the medium was changed to RPMI1640 with B-27 minus insulin and 5 μM IWP-2 (R&D Systems). On day 4, the medium was changed to RPMI1640 with B-27 minus insulin. On day 6, the medium was changed to RPMI1640 with B-27 serum-free supplement (Thermo Fisher), and on days 8 and 12, the same medium was exchanged to generate beating cardiomyocytes. We followed a previously reported protocol for metabolic selection of cardiomyocytes using lactate (*9*). On day 15, the cells were detached from the 12-well plate with 0.25% Trypsin-EDTA and resuspended in EB20 medium (Knockout DMEM (Thermo Fisher) containing 10% FBS, 1x GlutaMAX-I (Thermo Fisher), 100 μM MEM NEAA (Thermo Fisher), and 0.1 mM β-mercaptoethanol). The cell suspension was centrifuged at 300 × *g* for 5 min and the supernatant was removed. The cells were resuspended in RPMI1640 with B-27 serum-free supplement and 10 μM Y-27632 and plated into a Matrigel-coated well of a 6-well plate. On day 16, the medium was changed to RPMI1640 with B-27 serum-free supplement without 10 μM Y-27632. On day 20, the medium was changed to lactate medium (DMEM without glucose and sodium pyruvate (Thermo Fisher) with 1% MEM Non-Essential Amino Acids Solution (Thermo Fisher), Glutamax (Glutamax), and 4mM Sodium L-Lactate (Sigma)). Differentiated cardiomyocytes were selected by changing the fresh lactate medium on days 22 and 24. On day 26, the medium was replaced with RPMI1640 with B-27 serum-free medium. At day 28, purified cardiomyocytes were ready for the assay.

### Mitochondrial and genomic DNA copy number quantification by droplet digital PCR

We designed MT-ND4 and RPP30 specific primers and fluorescent hydrolysis probes to detect mitochondrial and genomic DNA using droplet digital PCR (ddPCR). The primer and probe sequences are shown in Table S4.

The 20x ddPCR assay premixtures were composed of forward and reverse primers (18 μM each) and MT-ND4 and RPP30 probes (5 μM each). We mixed the following reagents in a 96-well plate to make a 25-μl reaction: 12.5 μl of ddPCR Supermix for Probes (no dUTP) (Bio-Rad Laboratories #186-3024), 1.25 μl of 20x assay, 10 U of HindIII-HF (New England BioLabs #R3104S), 100–150 ng of genomic DNA in water, and water up to 25 μl. Droplets were generated with 20 μl of the premixed reaction and a QX100 Droplet Generator according to the manufacturer’s instructions (Bio-Rad Laboratories) and transferred to a 96-well PCR plate for standard PCR on a C1000 Thermal Cycler with a deep-well block (Bio-Rad Laboratories).

The thermal cycling program was as follows: (1) step 1, 95°C for 10 min; step 2, 94°C for 30 s; step 3, 59°C for 1 min; repeat steps 2 and 3 39 times; step 4, 98°C for 10 min, with all the steps ramped by 2°C/s. After PCR, the droplets were analyzed with a QX100 Droplet Reader (Bio-Rad Laboratories) with the “CNV2” option.

### Visualization and quantification of mitochondria in iPS-CMs

To stain the mitochondria and nuclei, the medium was changed to RPMI1640 with B27 containing 0.05 nM MitoTracker® Green FM probes (Thermo Fisher) and 1 μg/ml Hoechst 33342 (Nacalai Tesque) and incubated at 37°C for 15 min. The medium was then changed to RPMI1640 with B27, without MitoTracker and Hoechst 33342. The fluorescence of MitoTracker at 516 nm and Hoechst 33342 at 461 nm were observed using BZ-X810 fluorescence microscopy (Keyence). The area of mitochondria and number of nuclei were automatically calculated using the Fiji software program and a macro code modified based on previous reports^23,24^.

### Electron microscope

Cardiomyocytes were plated on chamber slides (Matsunami, SCS-N02). After culturing for 1 week, the cells were fixed with 4% paraformaldehyde (PFA) containing 0.05% calcium chloride in 0.1M cacodylate buffer, pH 7.3 (CB) for 30 min at 4°C, and fixed with 2% PFA, 2.5% glutaraldehyde, and 0.05% calcium chloride in 0.1M CB overnight. After rinsing 5 times in 4.5% sucrose in 0.1M CB, samples were postfixed with 2% osmium tetroxide in 0.1M CB for 2 h and dehydrated in a graded series of ethanol. Subsequently, the samples were flat embedded on glass chamber slides and polymerized in epoxy resin (EPON 812, TAAB) at 60°C for 48 h, then adhered to another block of epoxy resin and stained with toluidine blue to visualize the cells. Embedded samples were cut into ultrathin sections (50-80 nm thickness) using an ultramicrotome (PowerTomeX, RMC Boeckeler) and placed in formvar-coated single-slot grids. After staining with uranyl acetate and lead citrate, the ultrathin sections were observed under a transmission electron microscope (JEM-1400, JEOL) equipped with a bottom-mounted CCD camera.

### Measurement of impedance of iPS-CMs and terfenadine treatment

A total of 100,000 differentiated cardiomyocytes at day 28 were plated onto a well of NSP-96 Sensor Plates CardioExcyte96 (2.0 mm, Nanion Technologies) with RPMI1640 containing B-27 serum-free supplement and 10 μM Y-27632. The next day, the medium was changed to RPMI1640 with B-27 serum-free supplement without Y-27632 every day, and the cardiac impedance was recorded for 30 s at each recording points with 15 minutes interval over 1 h in total using CardioExcyte96 (Nanion Technologies) every day before changing the medium for 2 weeks. For terfenadine treatment on day 5 after seeding cardiomyocytes on the sensor plate, the cardiac impedance was recorded under normal conditions for 1 h. After that, terfenadine (Sigma) was added to the culture medium and the changes in cardiac impedance were recorded for 30 s at each point with 5 min intervals for 30 min in total. This analysis was conducted as previously reported^25^.

### Statistical analyses

All statistical analyses were performed using Microsoft^®^ Excel for Mac (ver. 16.94). Data are shown as the mean ± standard error of the mean (SEM) and individual data points with statistical significance. P-values were determined using an unpaired two-tailed Student’s t-test for comparison of two samples. In the figure legends, *n* indicates the number of biological replicates.

