## Supplementary figures S1-S6 for "Derivation of human post-mitotic cardiomyocytes from tetraploid iPSCs"

**The PDF file includes** Figures S1-S6.

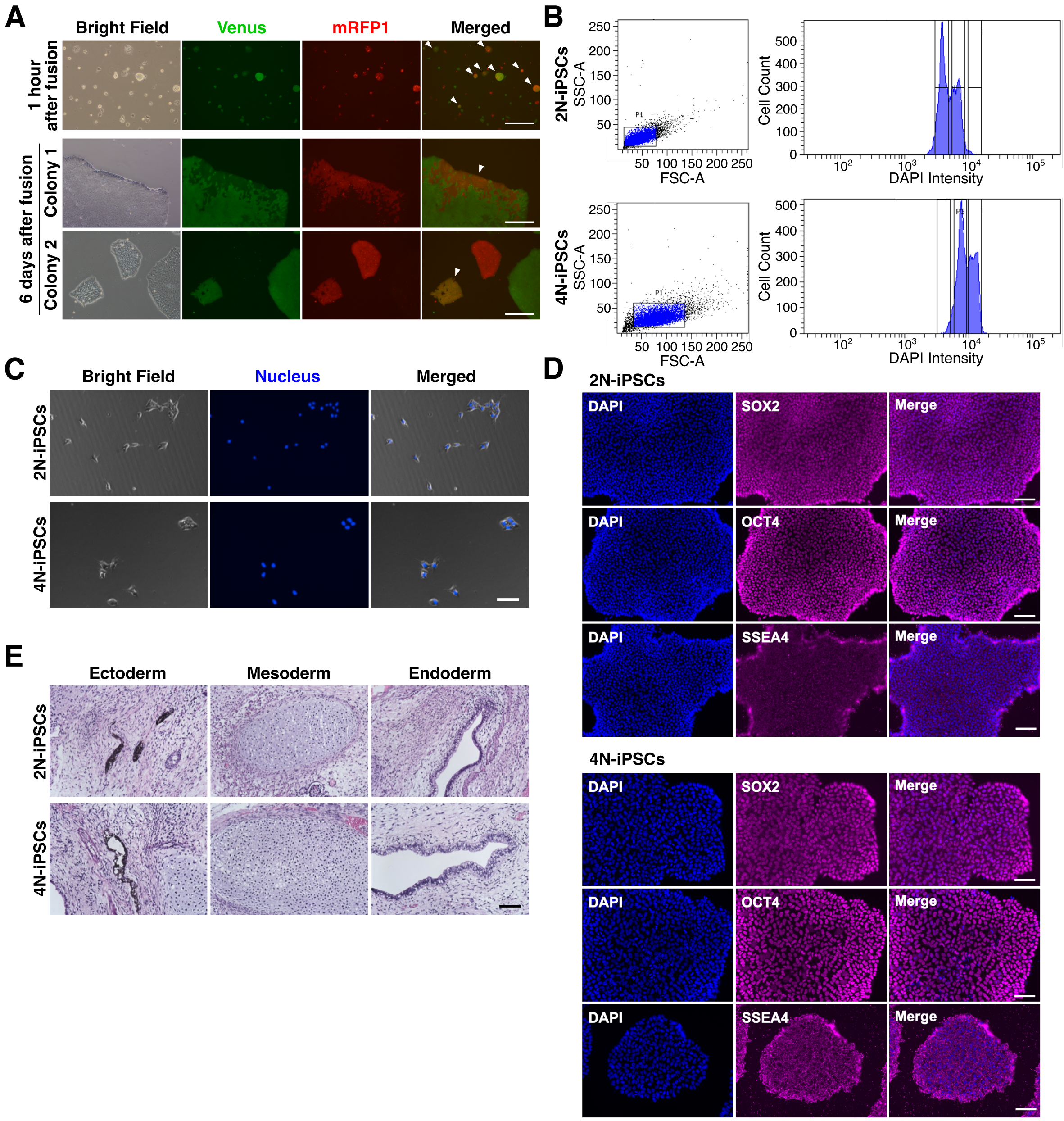


**Figure S1. Generation of 4N-iPSCs by fusion of 2N-iPSCs, related to Figure 1**

(A) Fluorescent images of fused iPSCs 1 hour and 6 days after cell fusion. Cells positive for both Venus and mRFP1 are indicated by white arrowheads. Scale bars, 200 μm.

(B) The cell cycle analysis of 2N- and 4N-iPSCs using flow cytometry to confirm a doubled DNA content in 4N-iPSCs.

(C) Nuclear staining in 2N- and 4N-iPSCs. Scale bars, 100 μm.

(D) Immunofluorescent staining of pluripotency markers, SOX2, OCT4, and SSEA4 in 2N- and 4N-iPSCs. Scale bars, 100 µm.

(E) Teratoma formation in NOD-SCID mice to confirm the pluripotency of 2N- and 4N-iPSCs. All three germ layers were observed. Scale bars, 125 µm.

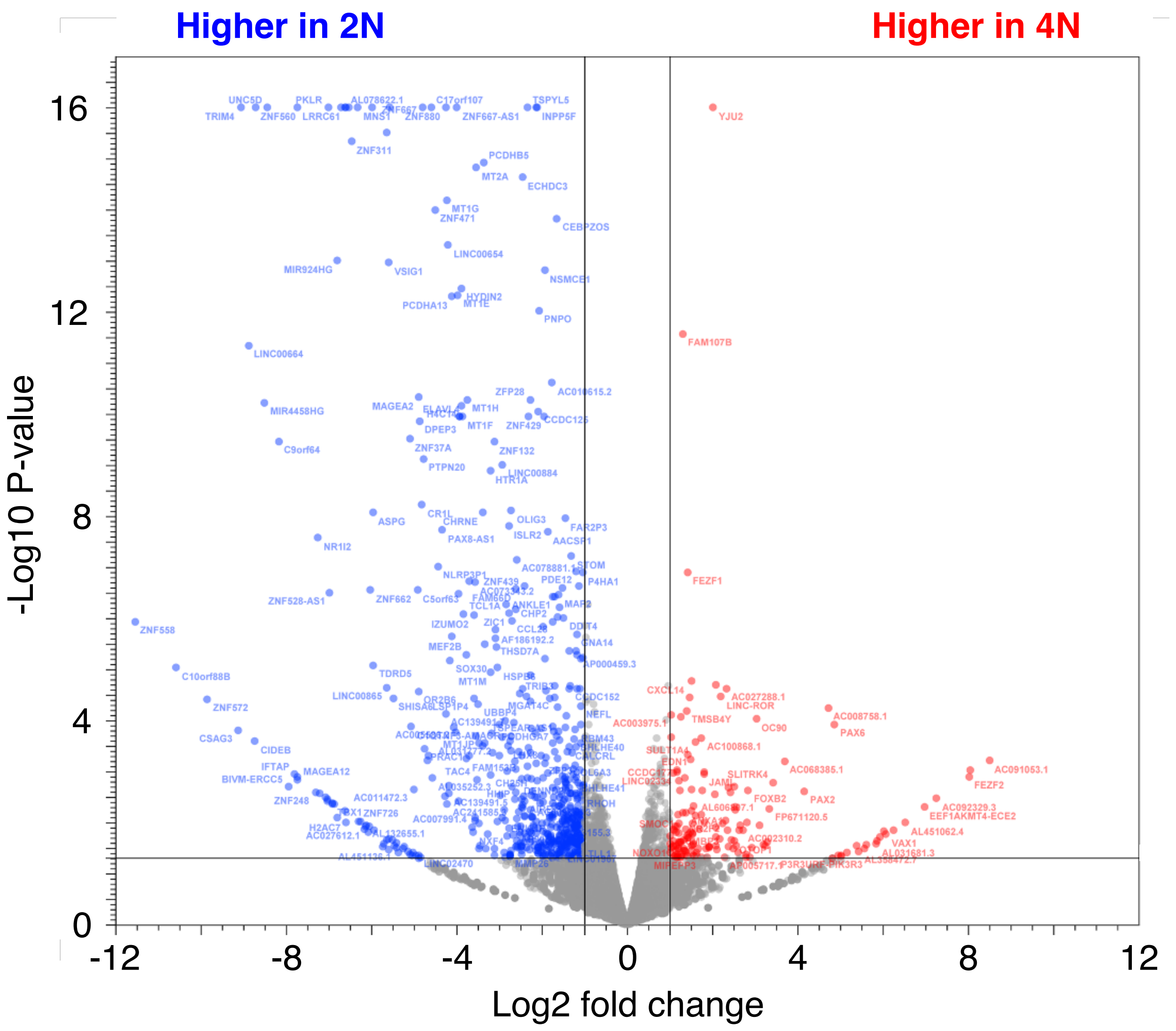


**Figure S2. Enhanced sensitivity of 4N-iPSCs to oxidative stress, related to Figure2**

Volcano plot to visualize differentially expressed genes (DEGs) between 2N- and 4N-iPSCs. Red and blue dots represent genes significantly upregulated or downregulated in 4N-iPSCs, respectively ($P<0.05$, $|\text{fold-change}|>2$).

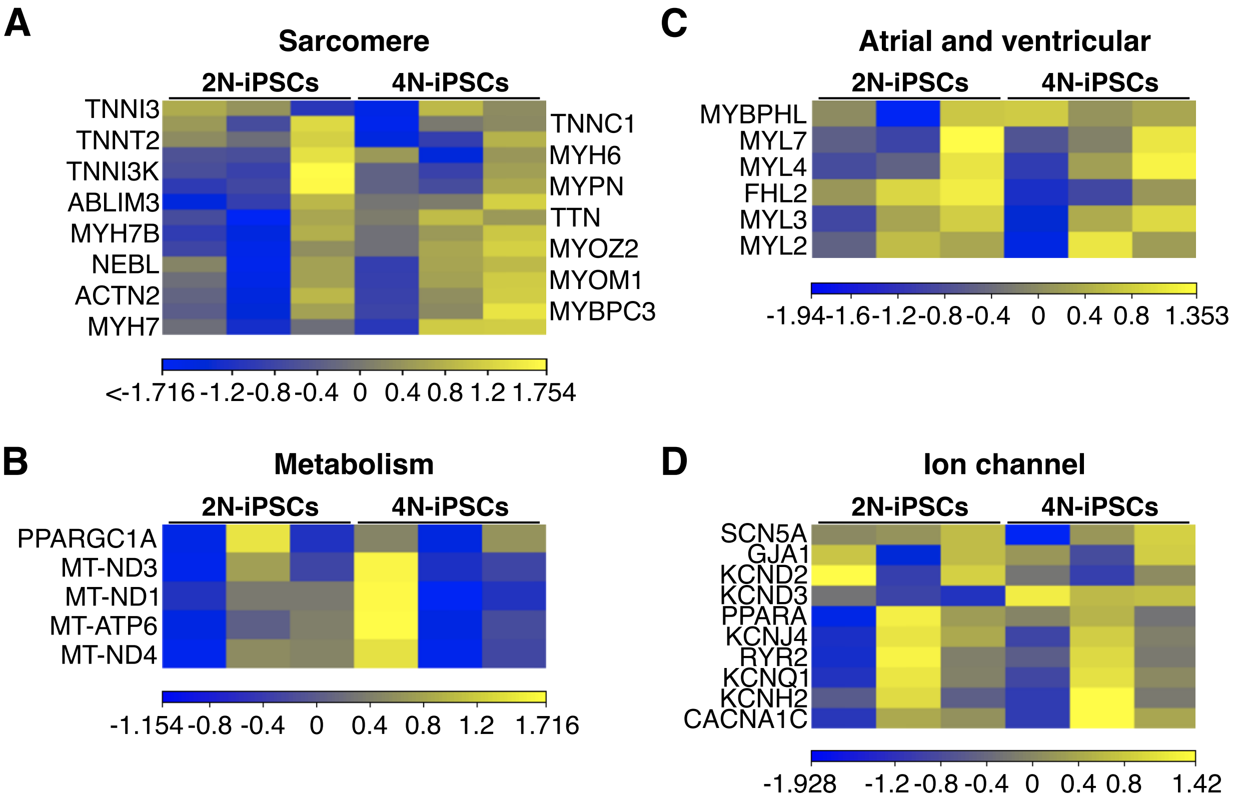


**Figure S3. The decreased expression of mitotic genes and increased mitochondria in 4N-iPS-CMs, related to Figure 3**

(A-D) Heatmap representation of the expression of genes involved in sarcomere structure (A), atrial and ventricular specification (B), Metabolism (C), and ion channel function (D) in 2N- and 4N-iPSCs.

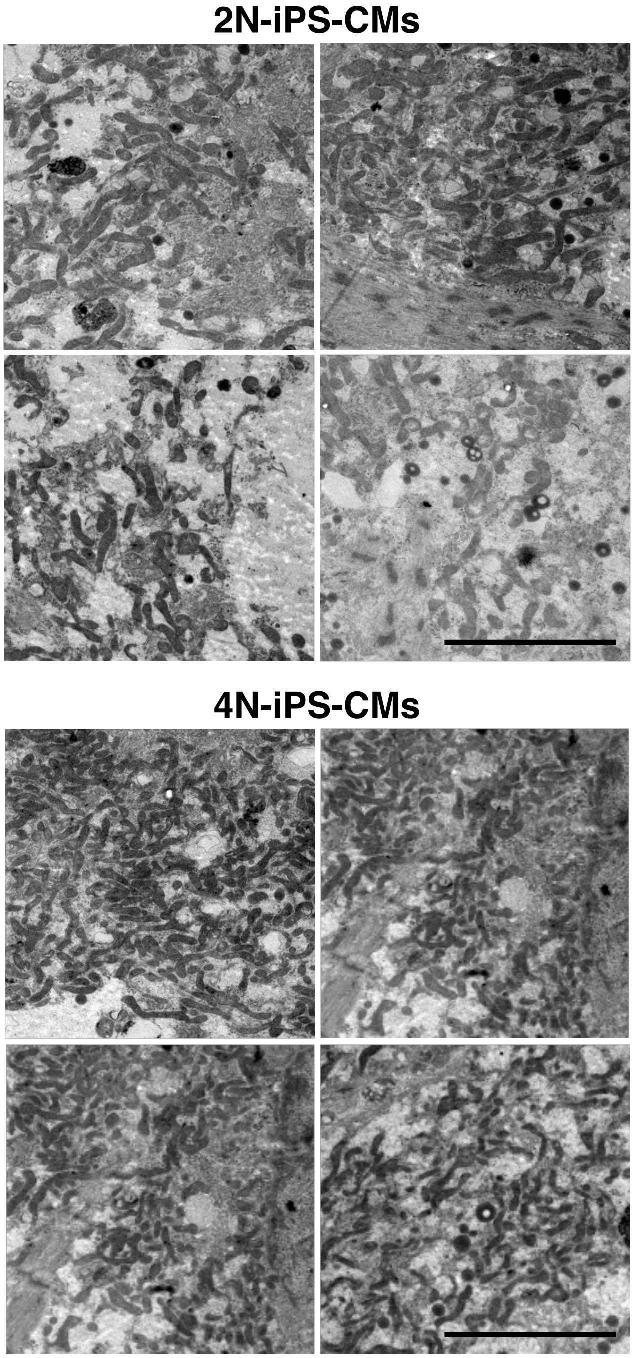


**Figure S4. The decreased expression of mitotic genes and increased mitochondria in 4N-iPS-CMs, related to Figure 3**

Electron microscopy images of mitochondria in 2N- (top) and 4N- (bottom) iPS-CMs. Scale bars, 5 μm.

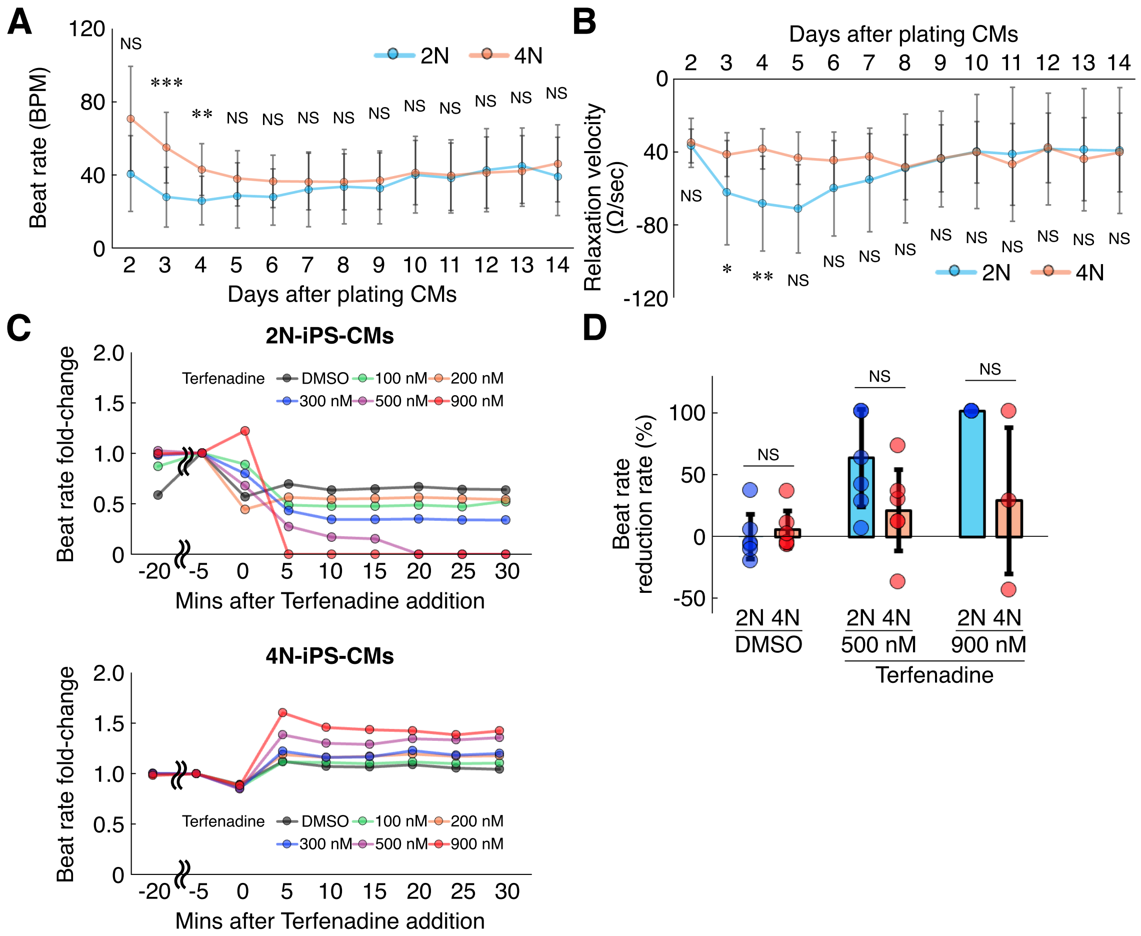


**Figure S5. Increased impedance and resistance to a potassium channel blocker in 4N-iPS-CMs, related to Figure 4**

(A and B) Beat rate (A) and relaxation velocity (B) of 2N- and 4N-iPS-CMs over two weeks after plating (*n*=15 for 2N-iPS-CMs, *n*=11 for 4N-iPS-CMs).

(C) Representative fold-changes of impedance amplitude of 2N- and 4N-iPS-CMs after the addition of different concentrations of Terfenadine.

(D) Beat rate reduction rate at 30 minutes after terfenadine treatment in 2N- and 4N-iPS-CMs (*n*=6 for DMSO and 500 nM, *n*=3 for 900 nM terfenadine).

Error bars, mean ± SEM: NS: not significant, *P*>0.05, **P*<0.05, ***P*<0.01, ****P*<0.001.

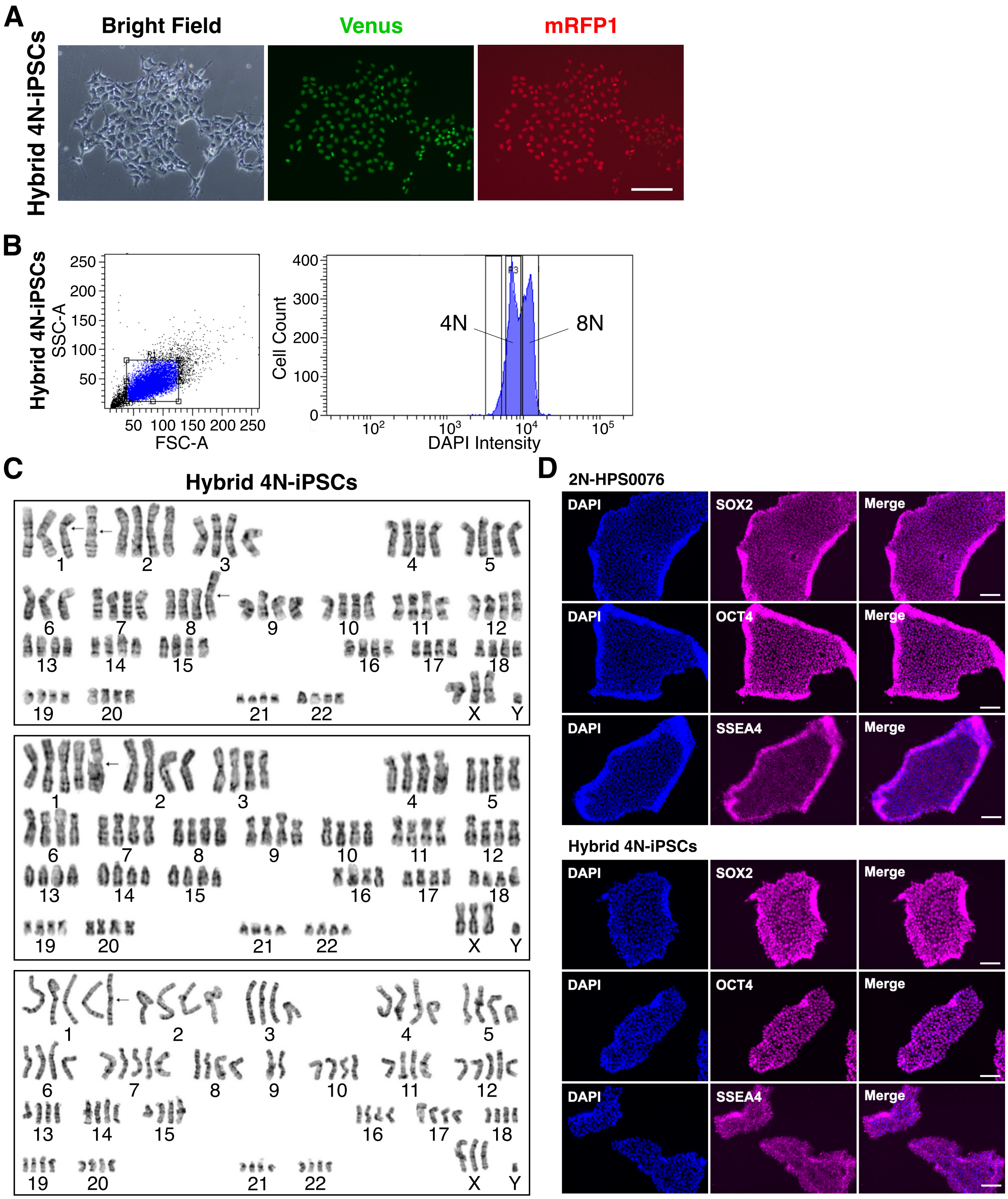


**Figure S6. Generation of 4N-iPSCs and 4N-iPS-CMs from two different individuals, related to Figure5**

(A) Fluorescent images of hybrid 4N-iPSCs. Scale bar, 200 μm.

(B) The cell cycle analysis of hybrid 4N-iPSCs using flow cytometry to confirm a doubled DNA content.

(C) The karyotype analysis of hybrid 4N-iPS cells. Chromosomal abnormalities were indicated by arrows.

(D) Immunofluorescent staining of pluripotency markers, SOX2, OCT4, and SSEA4 in diploid HPS0076 and hybrid 4N-iPSCs. Scale bars, 100 μm.
